## Supplemental material for "Polysaccharide breakdown products drive degradation-dispersal cycles of foraging bacteria through changes in metabolism and motility"

### SI Appendix for

**This PDF file includes:**

Figures S1 to S6
Table legends S1 to S17
Movie legends S1 to S5
Text

### SI Figures

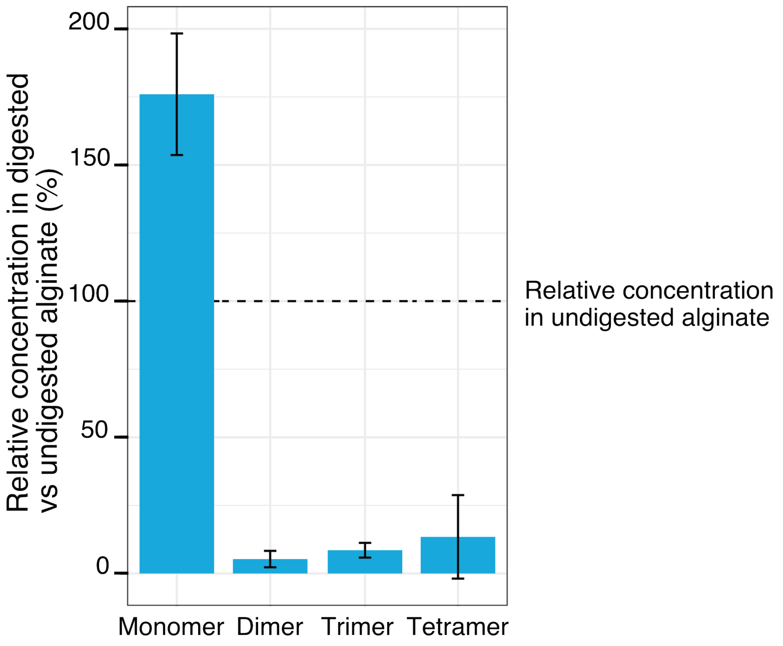


**Figure S1. Relative concentrations of the breakdown products of alginate after treatment with commercial alginate lyases.** LC-MS measurements of the digested and undigested alginate media, comparing the abundance of monomers, dimers, trimers, and tetramers in the digested alginate medium to the undigested one. The digestion was achieved by incubation with commercial alginate lyases for 48 hours. Bar heights depict the mean of four biological replicates whereas whiskers depict the standard deviation.

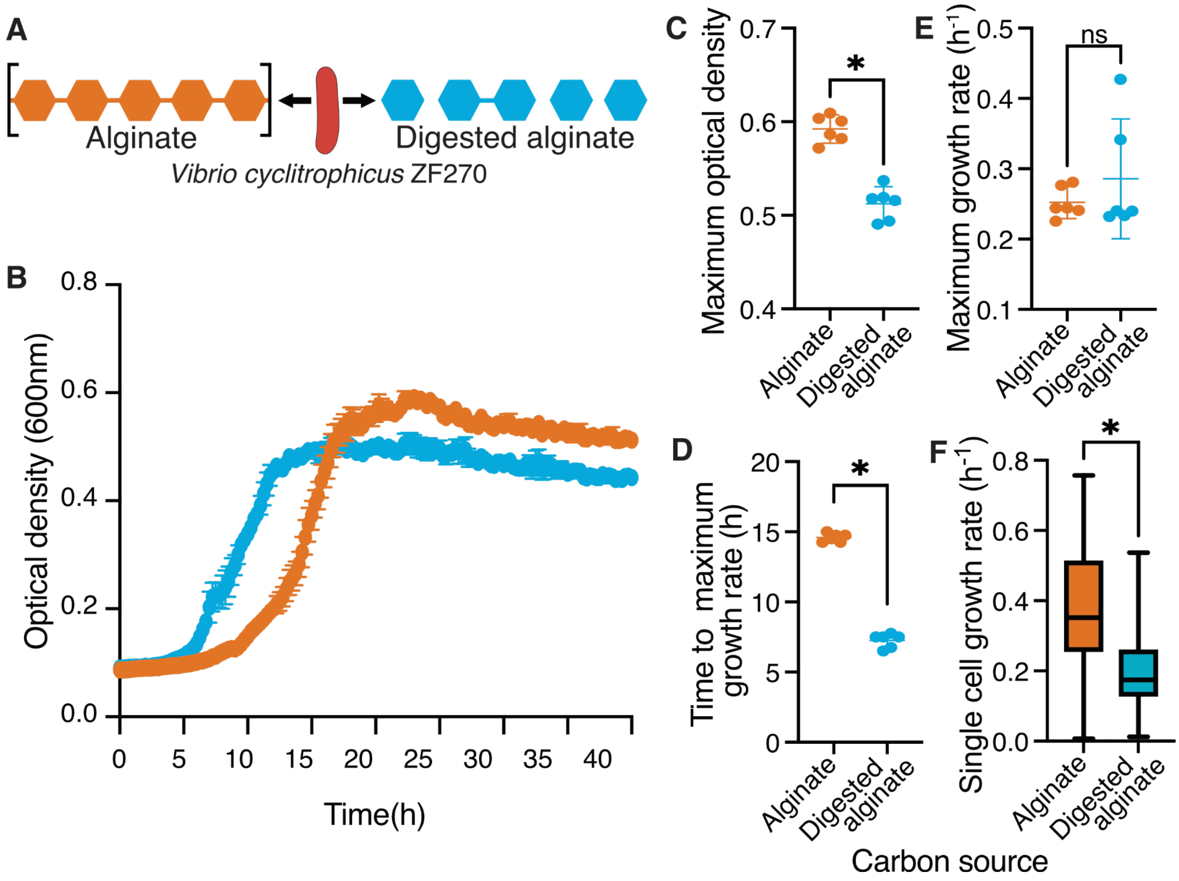


**Figure S2. Polymeric alginate increases lag times and yield of *Vibrio cyclitrophicus* ZF270 populations.** (A) *Vibrio cyclitrophicus* ZF270 was grown in microwell plates on 0.1 % (w/v) polymeric (alginate) or on 0.1% digested alginate as a sole carbon source. (B) Bacterial growth measured for 40 h using optical density (OD) at 600 nm. (C) Maximum OD600 on polymeric alginate (orange) or digested alginate (blue). Note that while the OD curves do not reach the exact same OD, plating cells on agar plates at 36 hours resulted in the same number of colonies (Fig. S3), indicating that the OD readings may be affected by the polysaccharide in the media. (D) Time to achieve maximal growth rates (lag time) and (E) maximum growth rates on polymeric alginate (orange) and digested alginate (blue). Circles indicate individual measurements whereas horizontal lines indicate the mean and whiskers the confidence intervals (CI) of six replicate populations. (F) Distribution of single cell growth rates of cells growing within microfluidic growth chambers on polymeric alginate (orange) or digested alginate (blue). Median growth rates were measured for each growth chamber for every 2h interval. Boxes extend from the 25th to 75th percentiles, whiskers indicate the 10th and 90th percentiles of median growth rates, and horizontal lines mark the median growth rates. Asterisks or ns indicate statistically significant or non-significant comparisons, respectively (independent samples t-test, in C: $p < 0.0001$, $t = 8.674$, $n = 6$; in D: $p = 0.34$, $t = 0.98$, $n = 6$; in E: $p < 0.0001$, $t = 30.73$, $n = 6$; in F: Mann-Whitney Test, $p < 0.0001$).

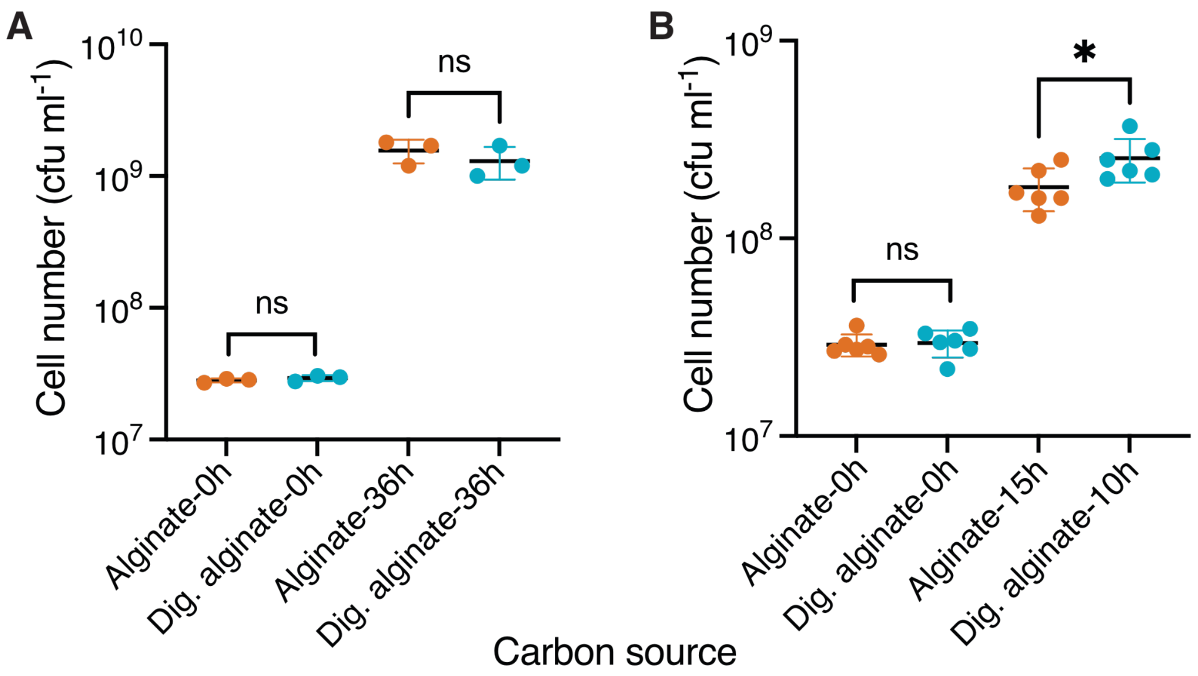


**Figure S3. Cell counts of *V. cyclotrophicus* ZF270 on alginate and digested alginate measured by plating assay.** (A) Cells were grown in shaking flasks and the yield was measured at the start of the experiment (0h) and after 36 h by plating on Marine Agar plates. Shown are colony forming units ml $^{-1}$ (cfu ml$^{-1}$) formed 24 h after plating. Circles indicate individual measurements whereas horizontal lines show the mean and whiskers represent the confidence intervals (CI) of 3 replicate populations. Cell numbers were statistically non-significant, depicted by *ns*, amongst groups (independent samples t-test, 0 h: $p = 0.33$, t = 1.093, n = 3; 36 h: $p = 0.39$, t = 0.95, n = 3). (B) Cells were grown on 0.1 % (w/v) polymeric or on 0.1% oligomeric alginate in shaking flasks and the yield was measured at the start of the experiment (0h) and when harvesting for RNA extraction, i.e., during exponential phase (based on Figure 1: 10 h for digested alginate (OD: 0.34) and 15 h for alginate (OD: 0.33)) by plating on Marine Agar plates. Shown are colony forming units ml $^{-1}$ (cfu ml$^{-1}$) formed 24 h after plating. Circles indicate individual measurements whereas horizontal lines show the mean and whiskers represent the confidence intervals (CI) of 3 replicate populations. Cell numbers were statistically non-significant, depicted by *ns*, amongst groups (independent samples t-test, 0 h: $p < 0.64$, $R^2 = 0.006$, t = 0.25, n = 6; 36 h: $p = 0.04$, $R^2 = 0.34$, t = 2.318, n = 3). Dig. alginate: digested alginate.

A

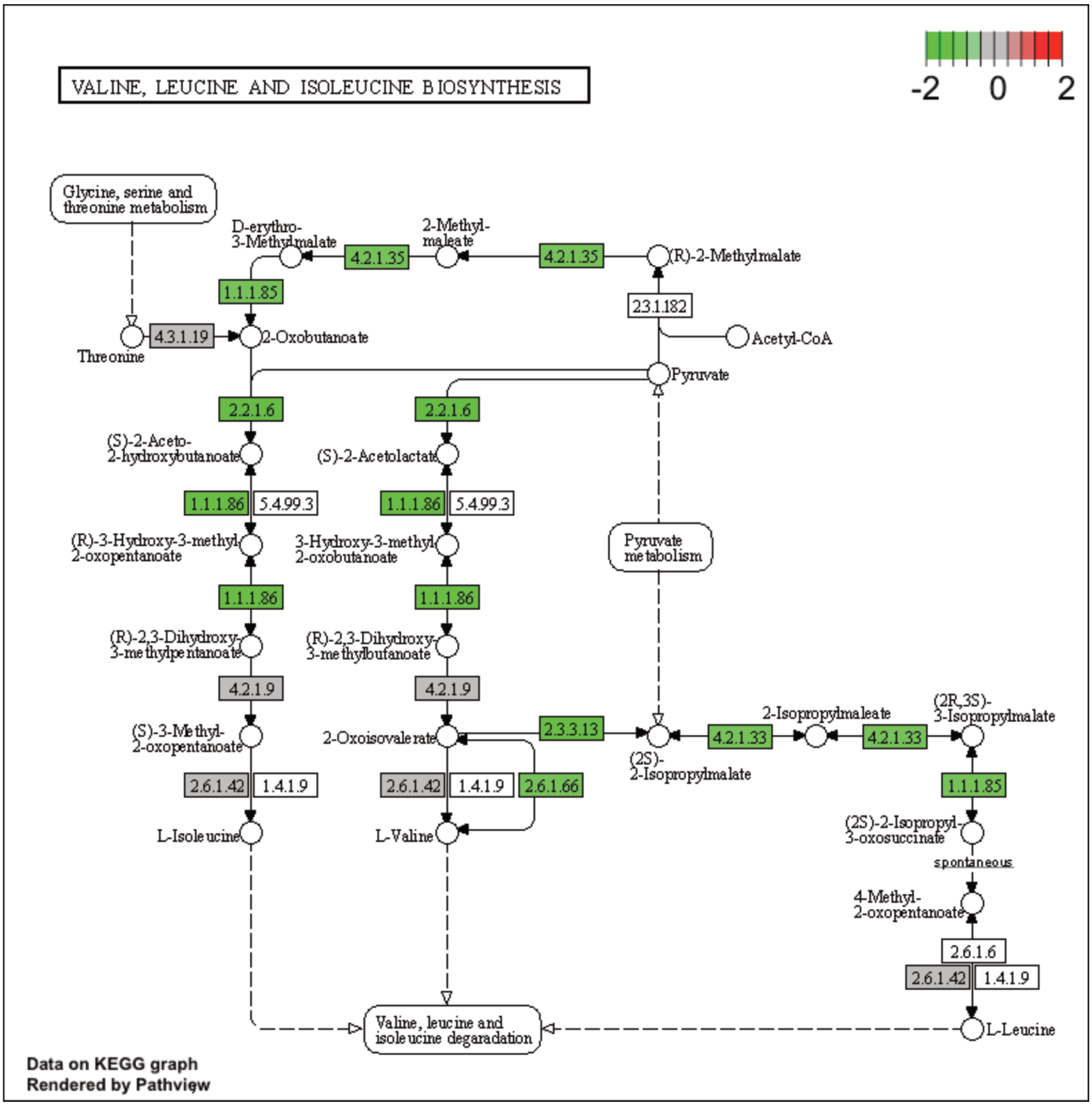

VALINE, LEUCINE AND ISOLEUCINE BIOSYNTHESIS
-2
0
2
Glycine, serine and threonine metabolism
D-erythro-3-Methylmalate
4.2.1.35
2-Methyl-maleate
4.2.1.35
(R)-2-Methylmalate
1.1.1.85
2.3.1.182
Threonine
4.3.1.19
2-Oxobutanoate
Acetyl-CoA
Pyruvate
2.2.1.6
2.2.1.6
(S)-2-Aceto-2-hydroxybutanoate
(S)-2-Acetolactate
1.1.1.86
5.4.99.3
1.1.1.86
5.4.99.3
(R)-3-Hydroxy-3-methyl-2-oxopentanoate
3-Hydroxy-3-methyl-2-oxobutanoate
Pyruvate metabolism
1.1.1.86
1.1.1.86
(R)-2,3-Dihydroxy-3-methylpentanoate
(R)-2,3-Dihydroxy-3-methylbutanoate
4.2.1.9
4.2.1.9
(S)-3-Methyl-2-oxopentanoate
2-Oxoisovalerate
2.3.3.13
(2S)-2-Isopropylmalate
4.2.1.33
2-Isopropylmaleate
4.2.1.33
(2R,3S)-3-Isopropylmalate
2.6.1.42
1.4.1.9
2.6.1.42
1.4.1.9
2.6.1.66
1.1.1.85
L-Isoleucine
L-Valine
(2S)-2-Isopropyl-3-oxosuccinate
spontaneous
4-Methyl-2-oxopentanoate
2.6.1.6
2.6.1.42
1.4.1.9
Valine, leucine and isoleucine degaradation
L-Leucine
Data on KEGG graph
Rendered by Pathview

**B**

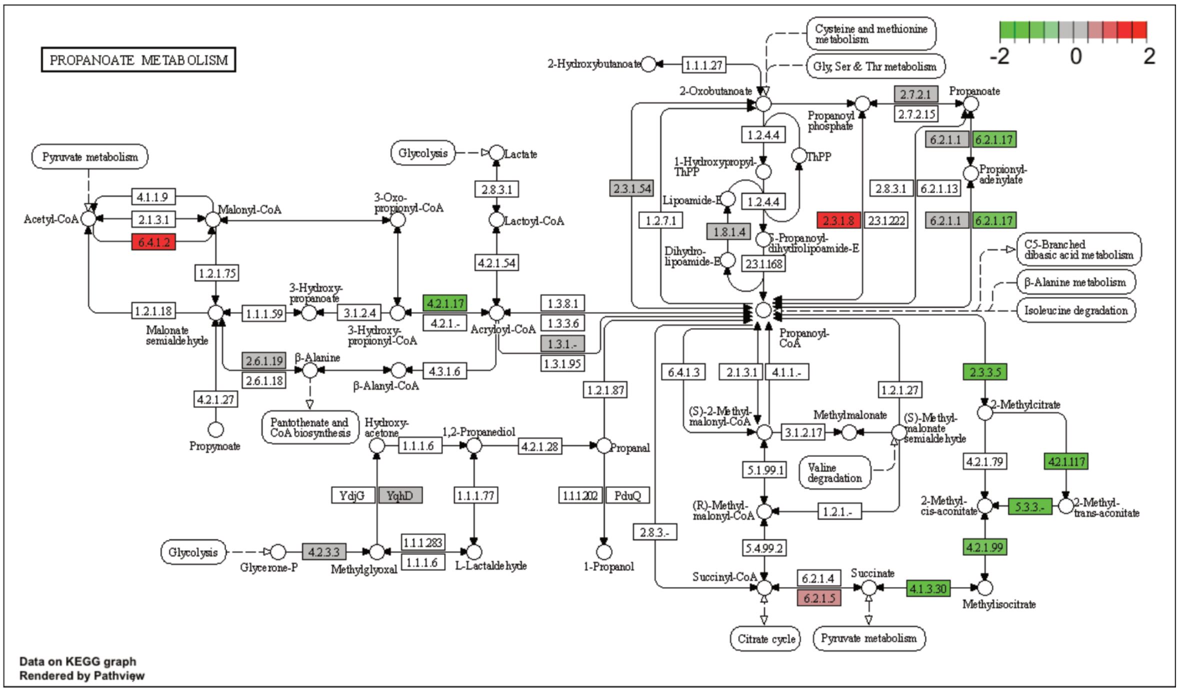
PROPANOATE METABOLISM
-2
0
2
Pyruvate metabolism
Acetyl-CoA
4.1.1.9
2.1.3.1
6.4.1.2
Malonyl-CoA
1.2.1.75
1.2.1.18
Malonate semialdehyde
4.2.1.27
Propynoate
3-Hydroxy-propanoate
1.1.1.59
3.1.2.4
3-Hydroxy-propionyl-CoA
3-Oxo-propionyl-CoA
2.6.1.19
2.6.1.18
β-Alanine
Pantothenate and CoA biosynthesis
β-Alanyl-CoA
4.3.1.6
Glycolysis
Lactate
2.8.3.1
Lactoyl-CoA
4.2.1.54
4.2.1.17
4.2.1.-
Acryloyl-CoA
1.3.8.1
1.3.3.6
1.3.1.-
1.3.1.95
Hydroxy-acetone
1.1.1.6
1,2-Propanediol
4.2.1.28
Propanal
1.2.1.87
YdjG
YqhD
1.1.1.77
1.1.1.202
PduQ
1-Propanol
Glycolysis
Glycerone-P
4.2.3.3
Methylglyoxal
1.1.1.283
1.1.1.6
L-Lactaldehyde
2-Hydroxybutanoate
1.1.1.27
Cysteine and methionine metabolism
Gly, Ser & Thr metabolism
2-Oxobutanoate
1.2.4.4
1-Hydroxypropyl-ThPP
ThPP
Lipoamide-E
1.2.4.4
1.8.1.4
S-Propanoyl-dihydrolipoamide-E
Dihydro-lipoamide-E
2.3.1.168
2.3.1.54
1.2.7.1
Propanoyl phosphate
2.7.2.1
2.7.2.15
Propanoate
6.2.1.1
6.2.1.17
Propionyl-adenylate
2.8.3.1
6.2.1.13
2.3.1.8
2.3.1.222
6.2.1.1
6.2.1.17
C5-Branched dibasic acid metabolism
β-Alanine metabolism
Isoleucine degradation
Propanoyl-CoA
6.4.1.3
2.1.3.1
4.1.1.-
1.2.1.27
(S)-2-Methyl-malonyl-CoA
3.1.2.17
Methylmalonate
(S)-Methyl-malonate semialdehyde
Valine degradation
5.1.99.1
(R)-Methyl-malonyl-CoA
1.2.1.-
5.4.99.2
2.8.3.-
Succinyl-CoA
6.2.1.4
6.2.1.5
Succinate
4.1.3.30
Methylisocitrate
Citrate cycle
Pyruvate metabolism
2.3.3.5
2-Methylcitrate
4.2.1.79
4.2.1.117
2-Methyl-cis-aconitate
5.3.3.-
2-Methyl-trans-aconitate
4.2.1.99
Data on KEGG graph
Rendered by Pathview

69

C

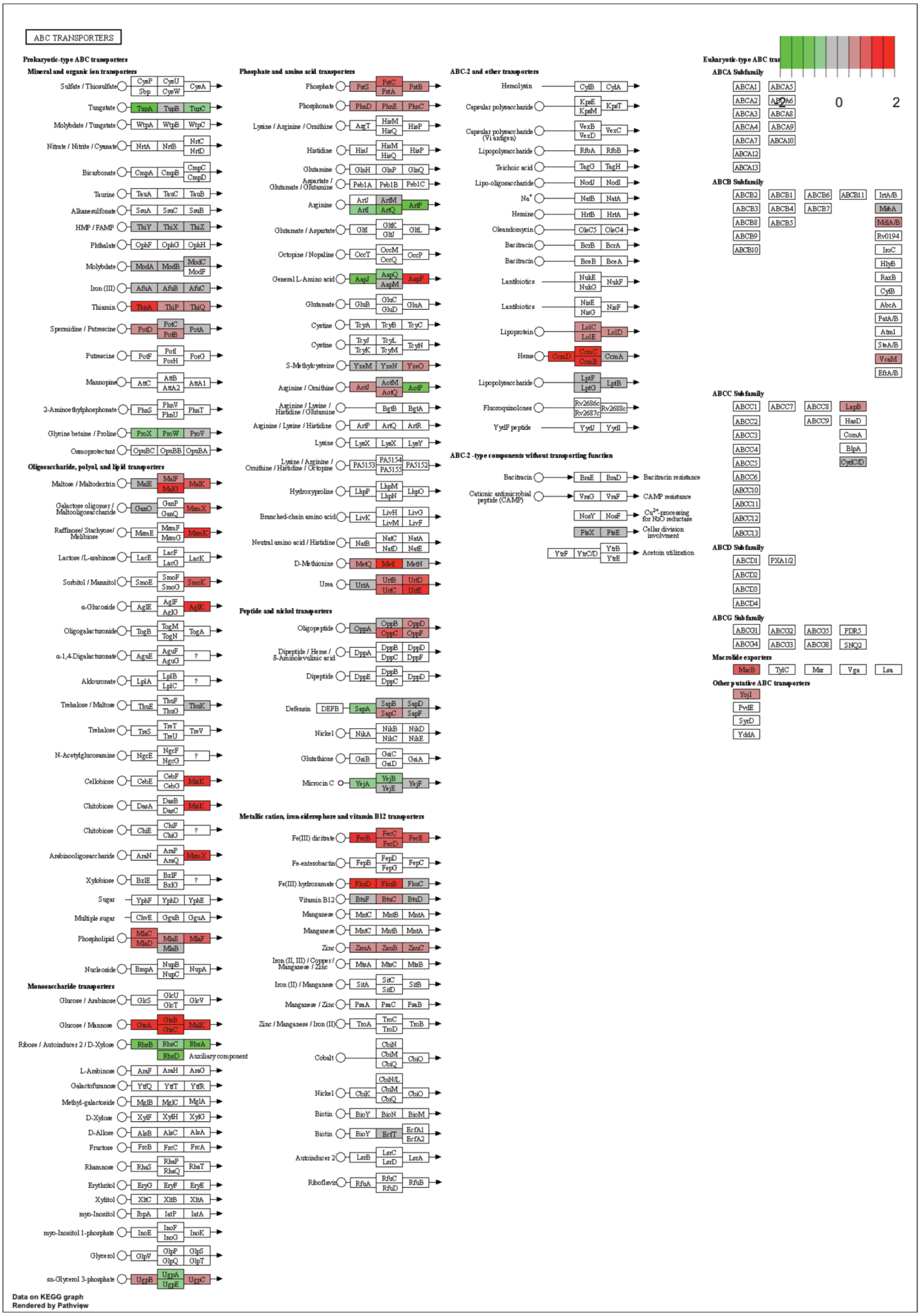

ABC TRANSPORTERS
Prokaryotic-type ABC transporters
Mineral and organic ion transporters
Phosphate and amino acid transporters
ABC-2 and other transporters
Eukaryotic-type ABC tra
-2
0
2
ABCA Subfamily
ABCB Subfamily
ABCC Subfamily
ABCD Subfamily
ABCG Subfamily
Macrolide exporters
Other putative ABC transporters
Oligosaccharide, polyol, and lipid transporters
Monosaccharide transporters
Peptide and nickel transporters
Metallic cation, iron-siderophore and vitamin B12 transporters
ABC-2-type components without transporting function
Data on KEGG graph
Rendered by Pathview

**D**

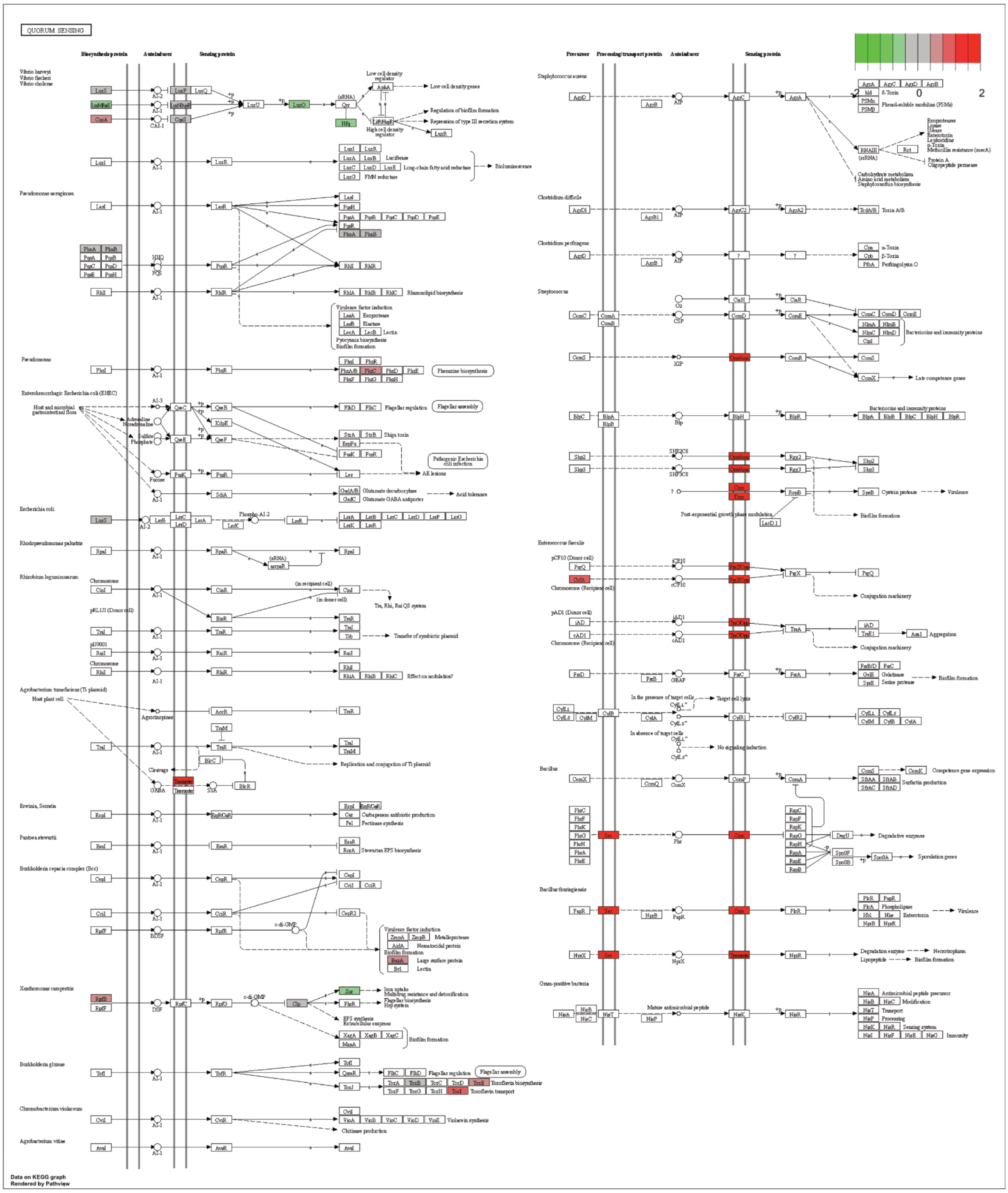
QUORUM SENSING
Biosynthesis protein
Autoinducer
Sensing protein
Precursor
Processing/transport protein
Staphylococcus aureus
Clostridium difficile
Clostridium perfringens
Streptococcus
Enterococcus faecalis
Bacillus
Bacillus thuringiensis
Gram-positive bacteria
Pseudomonas aeruginosa
Escherichia coli
Xanthomonas campestris
Burkholderia glumae
Chromobacterium violaceum
Agrobacterium vitiae
Bioluminescence
Biofilm formation
Virulence
Data on KEGG graph
Rendered by Pathview

**E**

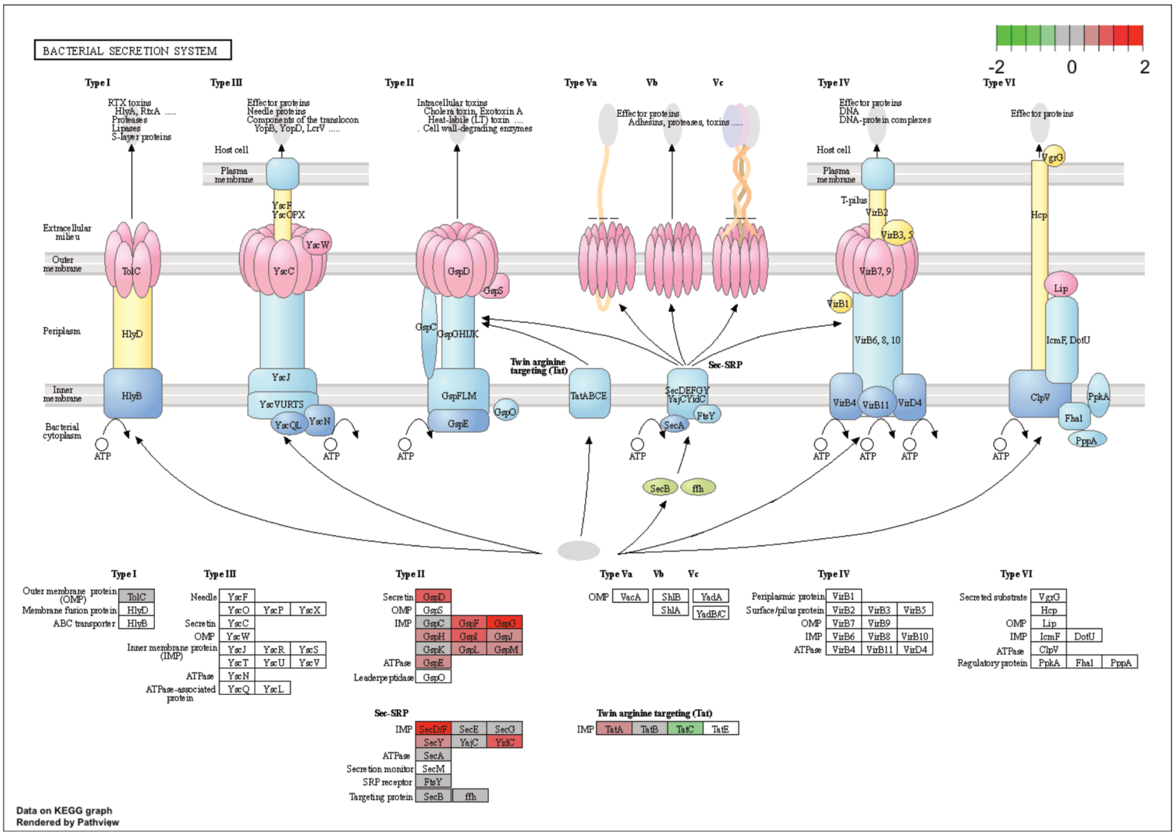
BACTERIAL SECRETION SYSTEM
-2 0 2
Type I
Type III
Type II
Type Va
Vb
Vc
Type IV
Type VI
RTX toxins
HlyA, RtxA .....
Proteases
Lipases
S-layer proteins
Effector proteins
Needle proteins
Components of the translocon
YopB, YopD, LcrV .....
Intracellular toxins
Cholera toxin, Exotoxin A
Heat-labile (LT) toxin .....
Cell wall-degrading enzymes
Effector proteins
Adhesins, proteases, toxins .....
Effector proteins
DNA
DNA-protein complexes
Effector proteins
Host cell
Plasma membrane
Extracellular milieu
Outer membrane
Periplasm
Inner membrane
Bacterial cytoplasm
TolC
HlyD
HlyB
YscF
YscOPX
YscW
YscC
YscJ
YscVURTS
YscQL
YscN
GspD
GspS
GspC
GspGHIJK
GspFLM
GspE
GspO
Twin arginine targeting (Tat)
TatABCE
Sec-SRP
SecDEFGY
YajC YidC
SecA
FtsY
SecB
ffh
T-pilus
VirB2
VirB3, 5
VirB7, 9
VirB1
VirB6, 8, 10
VirB4
VirB11
VirD4
VgrG
Hcp
Lip
IcmF, DotU
ClpV
PpkA
FhaI
PppA
ATP
Outer membrane protein (OMP)
Membrane fusion protein
ABC transporter
Needle
Secretin
OMP
Inner membrane protein (IMP)
ATPase
ATPase-associated protein
YscX
YscR
YscS
YscT
YscU
YscV
YscL
IMP
Leaderpeptidase
GspF
GspG
GspH
GspI
GspJ
GspK
GspL
GspM
VacA
ShlB
ShlA
YadA
YadB/C
Periplasmic protein
Surface/pilus protein
VirB8
VirB10
Secreted substrate
Regulatory protein
DotU
Secretion monitor
SRP receptor
Targeting protein
SecD/F
SecE
SecG
SecY
YajC
YidC
SecM
TatA
TatB
TatC
TatE
Data on KEGG graph
Rendered by Pathview

72

**F**

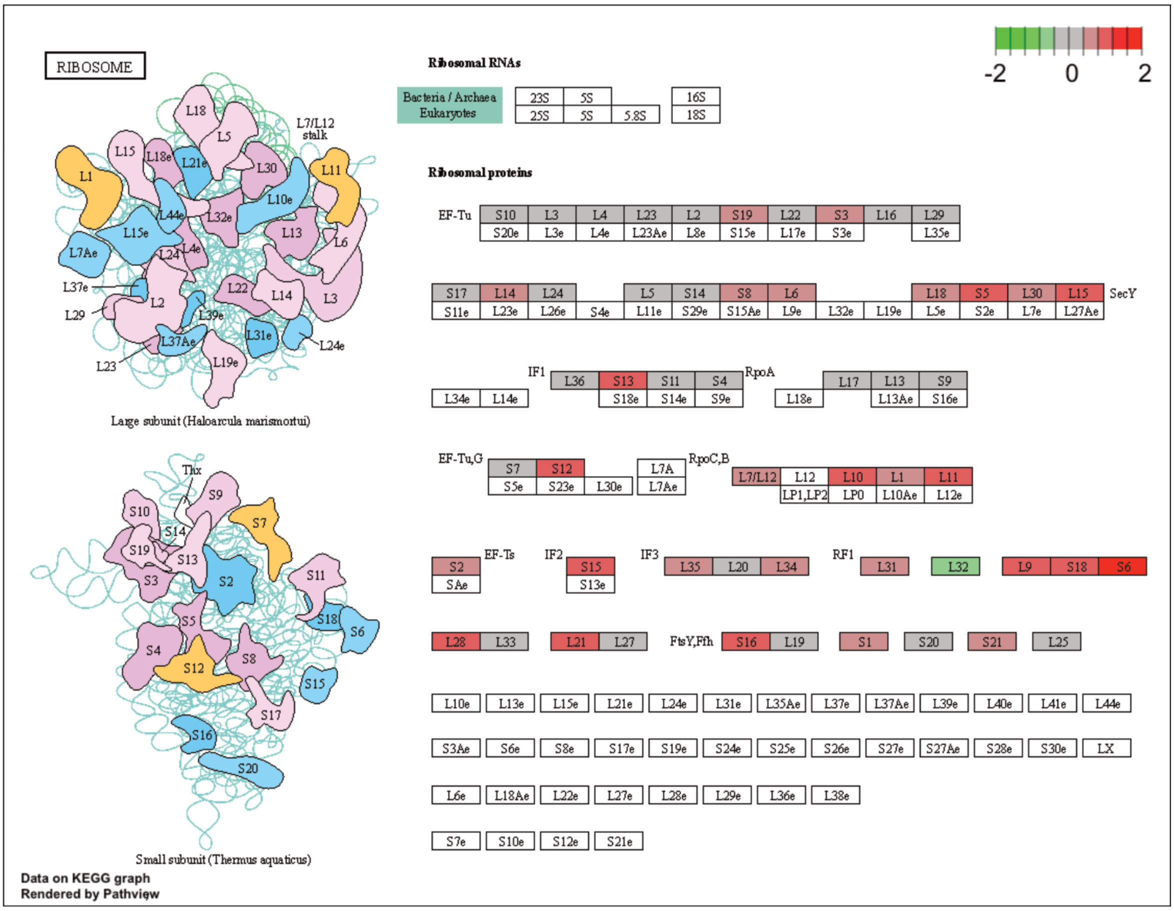
RIBOSOME
-2 0 2
Ribosomal RNAs
Bacteria / Archaea
Eukaryotes
23S 5S 16S
25S 5S 5.8S 18S
Ribosomal proteins
EF-Tu
S10 L3 L4 L23 L2 S19 L22 S3 L16 L29
S20e L3e L4e L23Ae L8e S15e L17e S3e L35e
S17 L14 L24 L5 S14 S8 L6 L18 S5 L30 L15
SecY
S11e L23e L26e S4e L11e S29e S15Ae L9e L32e L19e L5e S2e L7e L27Ae
IF1
RpoA
L34e L14e L36 S13 S11 S4 L17 L13 S9
S18e S14e S9e L18e L13Ae S16e
EF-Tu,G
S7 S12
S5e S23e L30e
L7A
L7Ae
RpoC,B
L7/L12 L12 L10 L1 L11
LP1,LP2 LP0 L10Ae L12e
S2 EF-Ts IF2 S15 IF3 L35 L20 L34 RF1 L31 L32 L9 S18 S6
SAe S13e
L28 L33 L21 L27 FtsY,Ffh S16 L19 S1 S20 S21 L25
L10e L13e L15e L21e L24e L31e L35Ae L37e L37Ae L39e L40e L41e L44e
S3Ae S6e S8e S17e S19e S24e S25e S26e S27e S27Ae S28e S30e LX
L6e L18Ae L22e L27e L28e L29e L36e L38e
S7e S10e S12e S21e
L7/L12 stalk
L1 L11 L18 L5 L15 L18e L21e L30 L10e L44e L32e L15e L13 L6 L7Ae L24 L4e L37e L29 L2 L22 L14 L3 L39e L31e L37Ae L24e L23 L19e
Large subunit (Haloarcula marismortui)
Thx
S10 S9 S14 S7 S19 S13 S3 S2 S11 S5 S18 S6 S4 S12 S8 S15 S17 S16 S20
Small subunit (Thermus aquaticus)
Data on KEGG graph
Rendered by Pathview

73

G

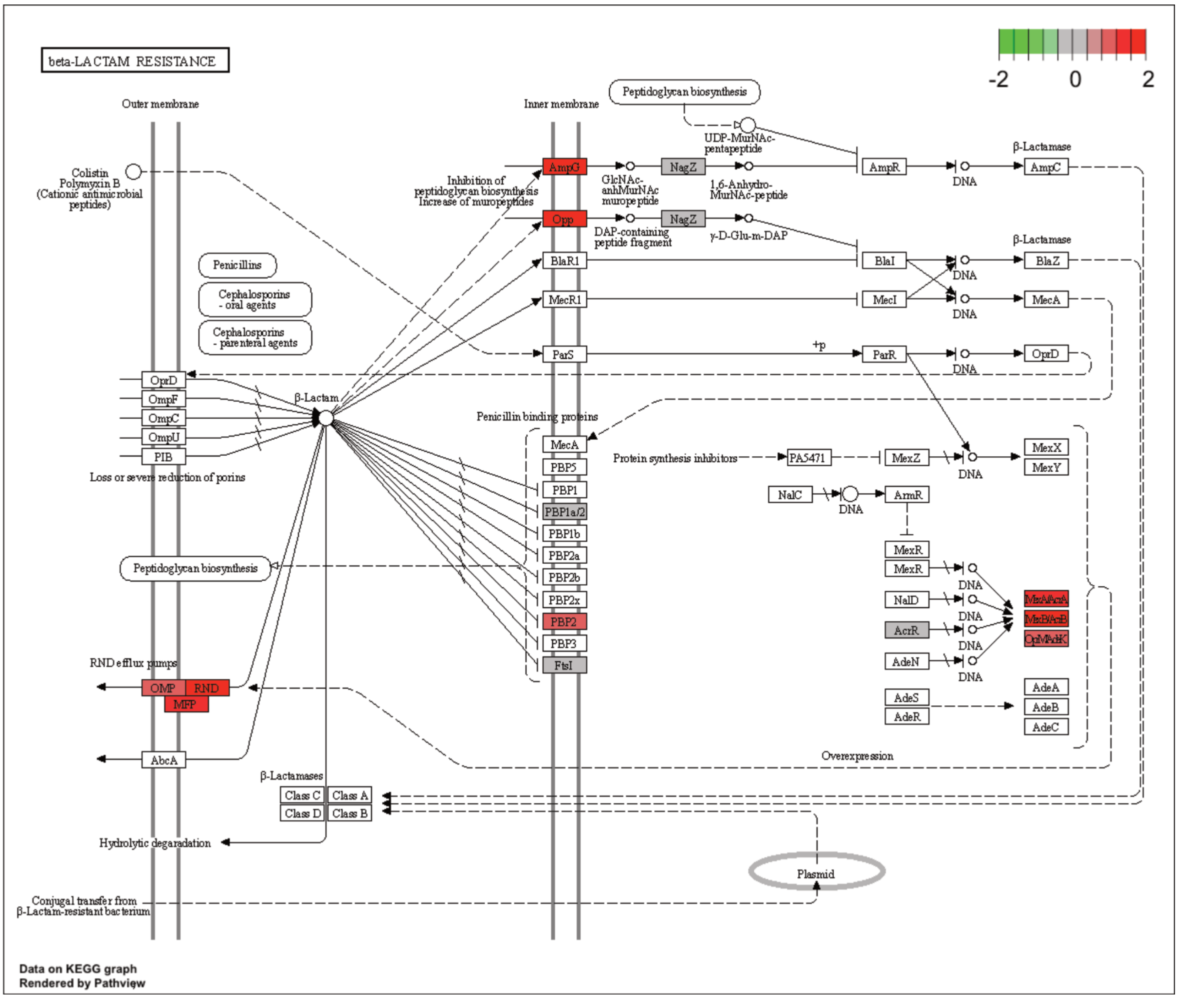


**Figure S4. KEGG maps of the significantly enriched KEGG pathways.** The labelled KEGG maps were created with the R package pathview v.1.35.0 for each KEGG pathway that was identified as a significantly enriched gene set by GSEA (see Materials and Methods). The color represents the log2 fold change of gene expression in *V. cyclitrophicus* ZF270 cells grown on digested alginate compared to alginate (green-grey-red color scale). No color (white) indicates that no gene of *V. cyclitrophicus* ZF270 could be mapped to this element of the KEGG map. Boxes represent gene products, circles represent metabolites.

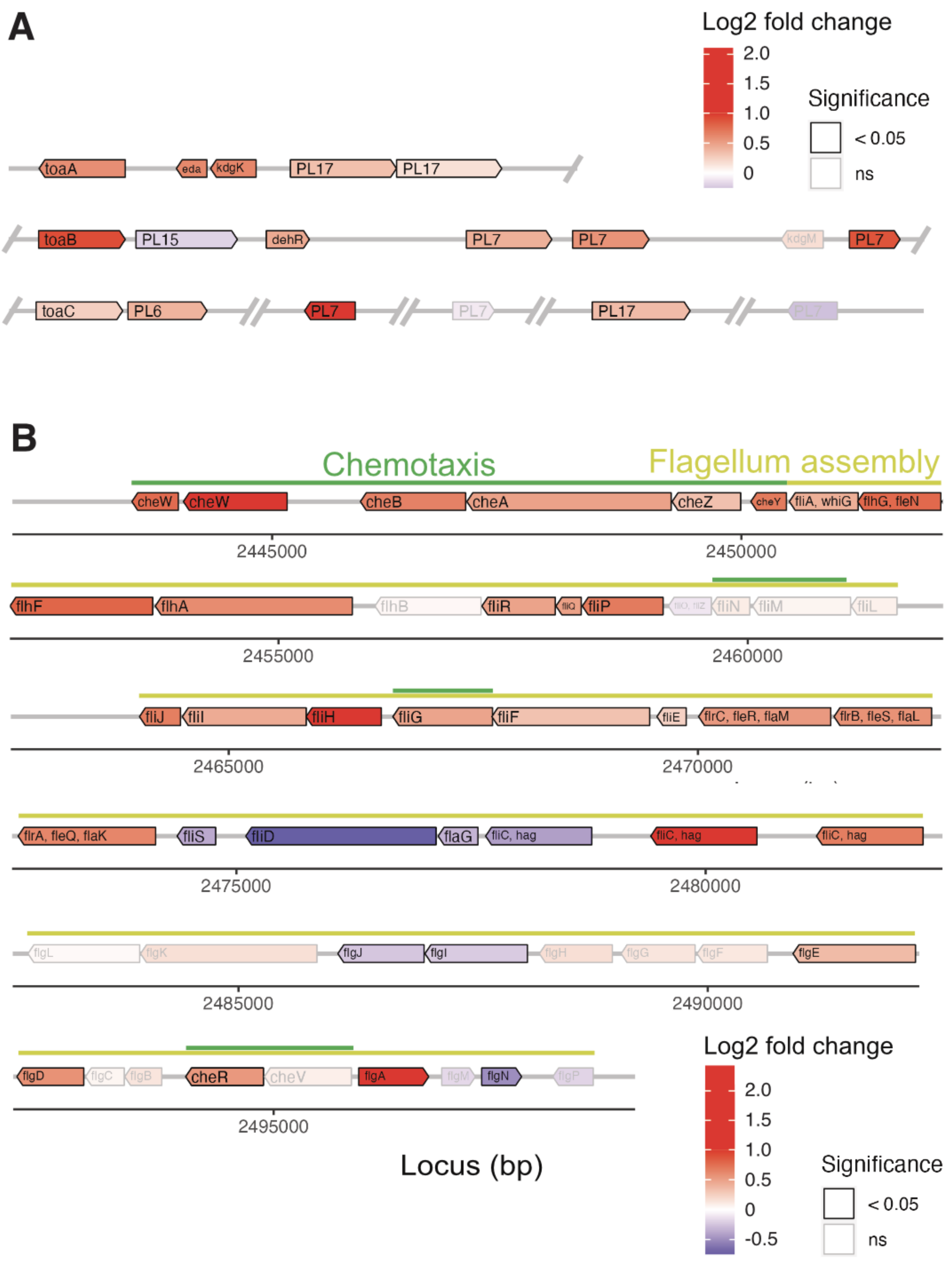


**Figure S5. Genomic location and differential expression of genes encoding (A) alginate catabolism and (B) the flagellum locus.** Gene expression of *V. cyclitrophicus* ZF270 on digested alginate was compared to the gene expression on alginate using genome-wide differential expression analysis. The log2 fold changes in gene expression of (A) alginate lyases (*PL6, PL7, PL15, PL17*), transporters (porin *kdgM*, symporter *toaB*, symporter *toaC*), and metabolic enzymes shunting into the ED pathway (DEHU reductase *DehR*, *kdgK*, *eda*), and (B) genes of the flagellum assembly locus and adjacent chemotaxis genes based on KEGG pathway 'Bacterial motility proteins' and 'Bacterial chemotaxis' is displayed. Differential expression analysis was performed with DESeq2 v1.32.0[56] to compute the Benjamini-Hochberg-adjusted Wald test *p*-value (box color) and log2 fold change (box fill) for each gene. For better visibility, genes that exhibited a log2 fold gene expression change greater than 1 (i.e., doubling of expression) or less than -1 (i.e., halving of expression) are designated maximum intensity of red or blue, respectively. In A, the location of the gene products was based on Wargacki et al. [17] with the exception of the alginate lyases (*PL6, PL7, PL15, PL17*) which were placed based on their signal peptide (S: extracellular, LS: membrane-anchored, none: cytosolic). OM: outer membrane; PM: periplasm; IM: inner membrane; PL: polysaccharide lyase family; *kdgM*: oligogalacturonate-specific outer membrane porin; *toaABC*: oligoalginate symporter; DEH: 4-deoxy-L-erythro-5-hexoseulose uronic acid; *dehR*: DEH reductase; KDG: 2-keto-3-deoxy-gluconate; *kdgK*: KDG kinase; KDPG: 2-keto-3-deoxy-6-phosphogluconate; *eda*: KDG-6-phosphate aldolase; GAP: glyceraldehyde 3-phosphate; ED: Entner-Doudoroff; ns: not significant.

### SI Table Legends

**Table S1. Differential gene expression of all genes of *V. cyclitrophicus* ZF270.** Genes of *V. cyclitrophicus* ZF270 were annotated by RASTtk. Differential expression analysis was performed on all genes with DESeq2 v1.32.0[53] to compute the log2 fold change in gene expression for each gene and the corresponding *p*-value by Benjamini-Hochberg-adjusted Wald test. Geneid: gene identifier of genome annotation file; Chr: chromosome; Start: start of gene in base pairs; End: end of gene in base pairs; Strand: DNA strand on which gene is located; Length: length of gene; L1_raw to L6_raw: raw read count of replicate 1 to 6 on digested alginate; P1_raw to P6_raw: raw read count of replicate 1 to 6 on polymeric alginate; L1_DESeq to L6_DESeq: DESeq2-normalized read count of replicate 1 to 6 on digested alginate; P1_DESeq to P6_DESeq: DESeq2-normalized read count of replicate 1 to 6 on polymeric alginate; baseMean: baseMean value computed with DESeq2; log2FoldChange: log2 fold change value computed with DESeq2 ; lfcSE: shrunken (posterior) standard deviation computed with DESeq2; stat: Wald statistic computed with DESeq2, i.e. the log2 fold change divided by lfcSE, which is compared to a standard Normal distribution to generate a two-tailed *p*-value; pvalue: Wald test *p*-value computed with DESeq2; padj: Benjamini-Hochberg-adjusted Wald test *p*-value computed with DESeq2; RASTtk_Annotation: gene annotation by RASTtk; RASTtk_Ontology_term: ontology term by RASTtk; BlastKOALA_KO: KEGG Orthology by BlastKOALA; BlastKOALA_KO_Definition: KEGG Orthology definition by BlastKOALA; BlastKOALA_KO_Score: weighted sum of BLAST bit scores computed by BlastKOALA; KEGG_pathway: ID of the KEGG category C associated with the KEGG Orthology (BlastKOALA_KO), i.e., KEGG pathway ID or KEGG BRITE ID; KEGG_pathway_descr: Description of the KEGG category C; KEGG_CategB: ID of the KEGG category B associated with the KEGG Orthology (BlastKOALA_KO); KEGG_CategA: ID of the KEGG category A associated with the KEGG Orthology (BlastKOALA_KO). All KEGG categories were based on https://www.kegg.jp/kegg-bin/show_brite?ko00001.keg , Mar 18 2021.

**Table S2. Genome-wide pathway enrichment analysis.** Performed on all gene sets of KEGG category C (KEGG pathways and KEGG BRITE categories) by Gene Set Enrichment Analysis algorithm (GSEA)[35]. Method: fgsea() function and described filtering (see Materials and Methods); KEGG_hierarchy: ID of KEGG category C; KEGG_entry: KEGG pathway or KEGG BRITE category; Description: Description of the KEGG category; pval: enrichment *p*-value of GSEA; padj: BH-adjusted *p*-value of GSEA; log2err: the expected error for the standard deviation of the *p*-value logarithm, ES: enrichment score, same as in Broad GSEA implementation; NES: normalized enrichment score, normalized to mean enrichment of random samples of the same size; size: size of gene set after removing genes not present in the genome of *V. cyclitrophicus* ZF270; Genes_total_ZF270: number of genes of *V. cyclitrophicus* ZF270 within the gene set, counting gene duplicates.

**Table S3 - S15 are subsets of Table S1 with the genes of each significantly enriched gene set.** "Valine, leucine and isoleucine biosynthesis (Table S3), "Propanoate metabolism (Table S4), "Ribosome (Table S5), "Secretion system (Table S6), "Bacterial secretion system (Table S7), "General secretion pathway" (Table S8), "Enzymes with EC numbers (Table S9), "Transporters (Table S10), "ABC transporters (Table S11), "Bacterial motility proteins (Table S12), "Quorum sensing (Table S13), Transcription factors (Table S14), "beta-Lactam resistance (Table S15).

**Table S16 and S17 are subsets of Table S1 with manually selected gene sets.** Differential expression of alginate lyases (*PL6, PL7, PL15, PL17*), transporters (porin *kdgM*, symporter *toaB*, symporter *toaC*), and metabolic enzymes shunting into the ED pathway (DEHU reductase *DehR*, *kdgK*, *eda*) (Table S16). Differential expression of genes of the flagellum locus, denoting association of each gene with the KEGG category of bacterial motility and chemotaxis (Table S17).

### SI Movie Legends

**Movie S1** Time-lapse video of *Vibrio cyclitrophicus* ZF270 cells within a representative microfluidics chamber fed with 0.1% alginate as the sole carbon source. Images were captured every 8 minutes. Cells are false colored based on the identity of their progenitor cells. Cells whose divisional history cannot be tracked are shown in blue.

**Movie S2** Time-lapse video of *Vibrio cyclitrophicus* ZF270 cells within a representative microfluidics chamber fed with 0.1% digested alginate as the sole carbon source. Images were captured every 8 minutes. Cells are false colored based on the identity of their progenitor cells. Cells whose divisional history cannot be tracked are shown in blue.

**Movie S3** Time-lapse video of *Vibrio cyclitrophicus* ZF270 cells within a representative microfluidics chamber fed with 0.1% alginate and then switched to 0.1% digested alginate as sole carbon sources. Timings of the switch and carbon sources are indicated on the images. Images were captured every 8 minutes.

**Movie S4** High frame rate (125 Hz, i.e., frames $s^{-1}$) time-lapse video of *Vibrio cyclitrophicus* ZF270 cells within a representative microfluidics chamber fed with 0.1% alginate as the sole carbon source.

**Movie S5** High frame rate (125 Hz, i.e., frames $s^{-1}$) time-lapse video of *Vibrio cyclitrophicus* ZF270 cells within a representative microfluidics chamber fed with 0.1% digested alginate as the sole carbon source.

### SI Text

**Protein amount of the alginate lyases added to create digested alginate**

Based on the following calculation, we conclude that the amount of protein added to the growth medium by the addition of alginate lyases is so small that we consider it negligible. In our experiment we used 1 unit/ml of alginate lyases in a 4.5 ml solution to digest the alginate. As the commercially purchased alginate lyases are 10,000 units/g, our 4.5 ml solution contains 0.45 mg of alginate lyase protein. The digested alginate solution diluted 45x when added to culture medium. This means that we added 0.18 µg alginate lyase protein to 1 ml of culture medium.

As a comparison, for 1ml of alginate medium, 1000µg of alginate is added or for 1 ml of Lysogeny broth (LB) culture medium, 3,500 µg of LB are added. Thus, the amount of alginate lyase protein that we added is ca. 5000 - 20,000 times smaller than the amount of alginate or LB that one would add to support cell growth. Therefore, we expect the growth that the digestion of the added alginate lyases would allow to be negligible.
